# Altered hypothalamic diffusivity in Parkinsonian autonomic dysfunction

**DOI:** 10.1101/767624

**Authors:** Célia Rousseau, Miriam Sklerov, Nina Browner, Yueh Z. Lee, Adrien Boucaud, Juan Prieto, Martin A. Styner, Eran Dayan

## Abstract

The pathophysiological basis of autonomic symptoms in Parkinson’s disease remains incompletely understood. The hypothalamus plays a key regulatory role in autonomic function and has been shown to be affected in Parkinson’s disease. Here, using diffusion magnetic resonance imaging, we investigated whether microstructural properties of the hypothalamus differ in Parkinson’s disease patients with high compared to low autonomic symptom burden.

Parkinson’s disease patients with low (n=25) and high (n=25) autonomic symptom burden were identified from a larger pool, based on scores from a questionnaire assessing autonomic symptoms in Parkinson’s disease (SCOPA-AUT). In each patient, we first segmented the hypothalamus manually, based on anatomical landmarks. Diffusivity measures were then extracted from the hypothalamus. Diffusivity measures calculated in the brainstem and the putamen were used to assess the specificity of the results.

Relative to patients with low autonomic symptom burden, patients with high burden showed increased mean, axial, and radial diffusivity in the hypothalamus. In contrast, we did not find significant group differences in any of these measures extracted from the brainstem or the putamen.

These results reveal consistent differences in the microstructural properties of the hypothalamus between patients with low and high autonomic symptom burden. Hypothalamic diffusivity properties can thus potentially be used as an imaging marker to assist in the identification of therapeutic targets for autonomic dysfunction in Parkinson’s disease.

## Introduction

The clinical diagnosis of Parkinson’s disease remains centered around the presence of motor symptoms. However, the disease is also associated with a range of non-motor symptoms, which often precede motor symptoms, and are frequent and disabling [1]. Autonomic dysfunction, in particular, is often reported early among Parkinson’s disease patients, and is known to have a detrimental effect on patients’ quality of life [1,2] and activities of daily living [3]. Furthermore, autonomic dysfunction portends a poorer overall prognosis in Parkinson’s disease [4,5]. Unfortunately, treatments for autonomic symptoms in Parkinson’s disease are limited. The mechanisms that underlie autonomic dysfunction in this disease, and in particular the contribution of the central nervous system [6], remain insufficiently understood.

Autonomic functions are controlled by the central autonomic network (CAN), a cortico-subcortical network composed of regions in the cerebral cortex, diencephalon and brainstem [7]. Within this network, the hypothalamus plays a major regulatory role, and is implicated in autonomic responses related to sleep-wake cycles, temperature, feeding, reproduction and stress [7,8]. The mixed neuronal populations within the hypothalamus are both directly and indirectly connected with hubs of the sympathetic and parasympathetic nervous systems [7]. Through intricate ascending and descending pathways, the hypothalamus is engaged in inflow and outflow of neuronal signals throughout the CAN [9]. This allows the hypothalamus to maintain the body’s homeostasis by adapting to internal and external stimuli [7,8].

The neuronal mechanisms underlying autonomic dysfunction in Parkinson’s disease remain incompletely understood. Postmortem studies have found Lewy bodies, the major pathological hallmark of Parkinson’s disease [10,11], in the hypothalamus [12,13]. It was recently shown that intrinsic functional interactions between the hypothalamus and the striatum and thalamus are reduced in Parkinson’s disease patients with high autonomic symptom burden [14]. It nevertheless remains unclear if these pathological findings and alterations in functional interactions reflect measurable changes in brain structure that can be detected in-vivo, as it cannot be assumed that modifications in functional connectivity translate to structural changes. Previous studies in Parkinson’s disease patients documented links between quantitative measures of autonomic function, primarily respiratory and cardiovascular, and macro and microstructural (diffusivity) brain properties [15,16]. However, it remains unknown if symptomatic burden is similarly associated with alterations in hypothalamic brain structure. In the present study, we set out to investigate whether the microstructural integrity of the hypothalamus, given its central regulatory role in the CAN and known pathologic involvement in Parkinson’s disease, is altered in Parkinson’s disease patients with high compared to low autonomic symptom burden.

## Methods

### Patients

All data were obtained from the Parkinson’s Progression Markers Initiative (PPMI) (www.ppmi-info.org/data), an observational multi-center study, where advanced imaging, biomarker, and standardized clinical and behavioral assessments were collected from a large group of de novo Parkinson’s disease patients. Full inclusion and exclusion criteria can be found online (www.ppmi-info.org). For each participating center, the Institutional Review Boards approved the study, and all participants provided written informed consent.

To select participants for analysis, we considered all Parkinson’s disease patients who had no missing diffusion MRI data obtained at visit one (as part of the protocol, two sequences were collected from the majority of subjects). Then, we sorted patients’ scores on the Scales for Outcomes in Parkinson’s disease-AUTonomic (SCOPA-AUT’, see next section) and selected the 25 Parkinson’s disease patients with the lowest scores (AUT_low_ group) and the 25 patients with the highest scores (AUT_high_ group). Two additional patients (one in each group) were excluded from analysis since their data failed automated quality control (QC, see bellow).

### Clinical assessments

Clinical assessments were administered as part of the PPMI study [17], evaluating motor and non-motor symptoms. For motor symptoms, we considered the Movement Disorder Society Unified Parkinson’s Disease Rating Scale (MDS-UPDRS) part III (Motor Examination). The Montreal Cognitive Assessment (MoCA) was used to evaluate cognitive function. The burden of autonomic symptoms was based on the SCOPA-AUT questionnaire [18], a reliable and valid [19] self-administered questionnaire addressing gastrointestinal (seven questions), urinary (six questions), cardiovascular (three questions), thermoregulatory (four questions), pupillomotor (one question) and sexual (two or three questions, depending on sex) symptoms. Total scores were calculated for each patient in each of the three scales. All scale data was acquired either at the same visit as the scans, or at the preceding visit.

### MRI data acquisition

MRI data were acquired on Siemens 3T Trio or Verio scanners (46/4 subjects, respectively) using an identical scanning protocol. We confirmed that group differences reported in the manuscript were retained when controlling for scanner type. 3D T1-weighted structural data were acquired with a sagittal magnetization-prepared rapid gradient echo sequence (MPRAGE) with GRAPPA (Flip angle = 9.0 deg; Field of view = 240×256×176 pixels^3^; Voxel size = 1×1×1 mm^3^; TE = 2.98 ms; TI = 900.0 ms; TR = 2300.0 ms; Pulse Sequence = GR/IR). Cardiac-gated diffusion-MRI data were acquired with 64 unique gradient directions (Field Strength = 3.0 T; Flip Angle = 90.0 deg, Field of view = 1044×1044×65 pixels^3^; Voxel size = 1.98×1.98×2.00 mm^3^; TE = 88.0 ms; TR = 900.0 ms; b-value = 1000.0 s/mm^2^, cardiac-triggered; Pulse Sequence = EP).

### Data analysis

#### Diffusion MRI data analysis

We first merged two successive diffusion weighted image (DWI) acquisitions (acquired during the same imaging session) into a single combined dataset. The merged diffusion data were quality controlled and corrected with *DTIPrep* [20] (https://www.nitrc.org/projects/dtiprep/), an open-source tool for identification and correction of common diffusion-MRI artifacts. Default settings were used in this automated QC preprocessing step. First, corrupted DWI data were identified and excluded, then baseline images were co-registered and averaged, to generate an enhanced average baseline image. All diffusion data were then registered to this average baseline and thereby corrected for Eddy-current and motion artifacts [20]. Subsequently, the standard tensor images (DTI) and their DTI property maps were computed, including mean diffusivity (MD), axial diffusivity (AD), radial diffusivity (RD) and fractional anisotropy (FA) maps. Tensor reconstruction was computed via weighted linear square fitting. Finally, the corrected DWI and DTI images were visually inspected.

#### Segmentation of the Regions of Interest

Regions of interest (ROIs) in the hypothalamus, the putamen and the brainstem were segmented for further analysis. The hypothalamus was manually segmented in each subject’s native structural (T1-weighted) space by a trained investigator, blinded to the clinical diagnosis at the time of segmentation via *ITK-SNAP* (http://www.itksnap.org). In each slice, the borders of the hypothalamus were first defined in the coronal view, based on anatomical landmarks [21]. In short, the segmentation began in the coronal section containing the anterior commissure and the optic chiasm. Anteriorly, the hypothalamus was localized below the anterior commissure, above the optic chiasm, demarcated laterally by the substantia innominata. Progressing posteriorly, the segmented hypothalamus was identified above the infundibulum, below the anterior thalamus, demarcated laterally by the substantia nigra. Then, in all axial sections, the outlines of the segmentation were smoothed, to refine sharp borders. ROIs in the bilateral putamen and the brainstem were segmented automatically in native individual-subject structural space using *FreeSurfer* (http://surfer.nmr.mgh.harvard.edu/). All segmentations underwent visual QC.

#### Co-registration and Metric extraction

The structural data were co-registered into the diffusion space via a two-step process. First, a rigid registration computed with the *BRAINSFit* module in *3D Slicer* (http://www.slicer.org), mapped the T1-weighted images into the DTI-AD maps. This rigid registration was then used to initiate a multimodal non-linear registration with Advanced Normalization Tools (*ANTs*, http://stnava.github.io/ANTs/), deformably mapping the T1-weighted images into the FA and B0 maps. Rigid and non-linear transformations were concatenated to produce global displacement fields that register the T1 structural images to the diffusion-MRI image space. The global deformation fields were also applied to the ROI segmentations of the hypothalamus, the brainstem and the putamen. Finally, within each co-registered ROI, the mean intensity of the diffusivity measures (MD, AD and RD) and FA, were extracted.

### Statistical analysis

Statistical analysis was performed using Statistical Package for Social Sciences Version 9 (*SPSS*, Inc., Chicago IL). To compare the two groups of Parkinson’s disease patients, t-tests were applied (2-tailed; homogeneity of variances assumed) and the significance level was fixed at p<0.05. We additionally controlled for clinical variables, which were significantly different between the AUT_low_ and AUT_high_ groups by performing an analysis of covariance (ANCOVA).

## Results

The aim of the study was to test whether Parkinson’s disease patients with high autonomic symptom burden exhibit microstructural alterations in the hypothalamus, a key structure of the CAN, relative to patients with low autonomic symptom burden.

### Demographic and clinical characteristics

Twenty-five Parkinson’s disease patients with low autonomic symptom burden (AUT_low_ group, 10 women, mean age = 57.4 ± 1.77 years) and 25 Parkinson’s disease patients with high symptom burden (AUT_high_ group, 9 women, mean age = 62.5 ± 1.67 years) were included in the study (Table 1). As expected, total SCOPA-AUT scores were significantly different (Fig. 1A) between the two groups (t_48_ = 11.782, p < 0.001). The AUT_high_ group was also older compared to the AUT_low_ group (t_48_ = −2.095, p = 0.042). In contrast, disease duration, MoCA scores and MDS-UPDRS Part III scores were not significantly different between the two groups (disease duration: t_48_ = −0.588, p = 0.560; MoCA scores: t_48_ = 1.227, p = 0.226; MDS-UPDRS Part III scores: t_48_ = −1.765, p = 0.084). Of note, disease duration was calculated based on the time from Parkinson’s disease diagnosis to diffusion MRI data acquisition. We also assessed disease duration relative to the approximate date of symptom onset (AUT_low_ group, disease duration = 28.40 ± 27.74 months; AUT_high_ group, disease duration = 30.49 ± 46.03 months), and found no significant differences between the two groups (t_48_ = −0.255, p = 0.8).

**Table 1:**
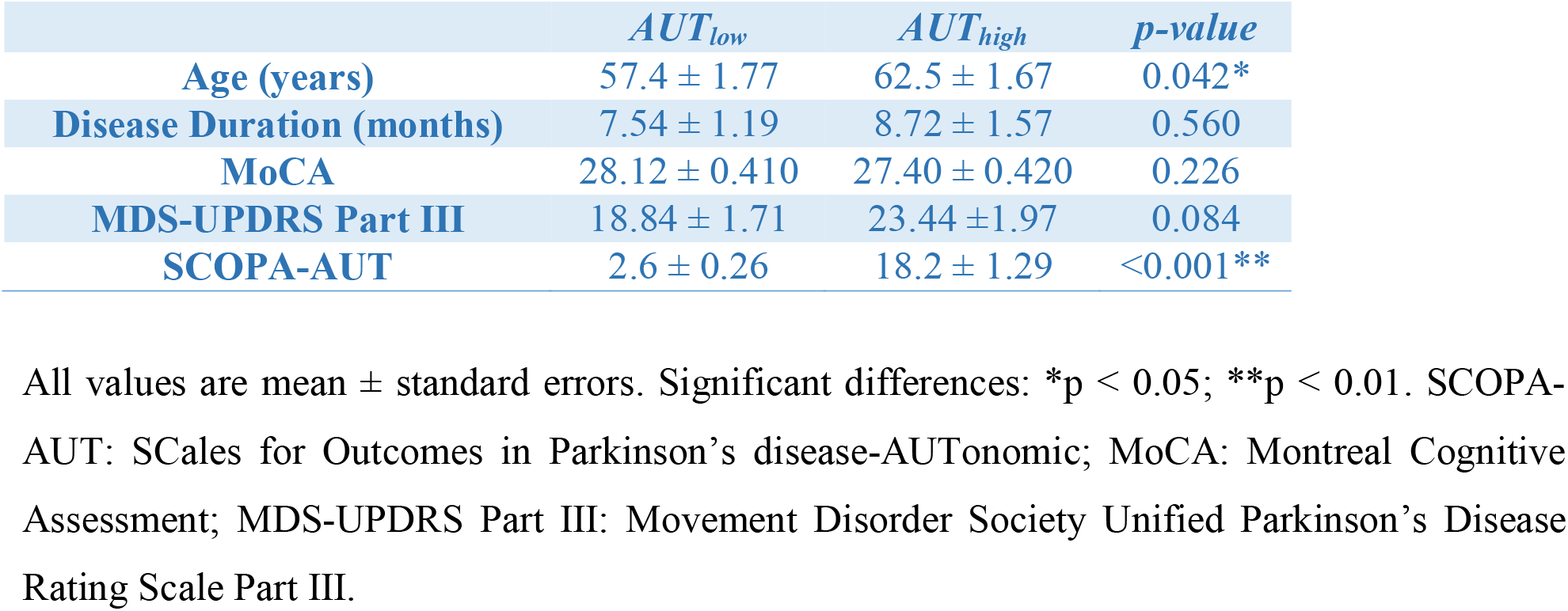
Demographic and clinical characteristics of Parkinson’s disease patients in the low (AUT_low_) and high (AUT_high_) autonomic symptom burden groups.

**Figure 1:**
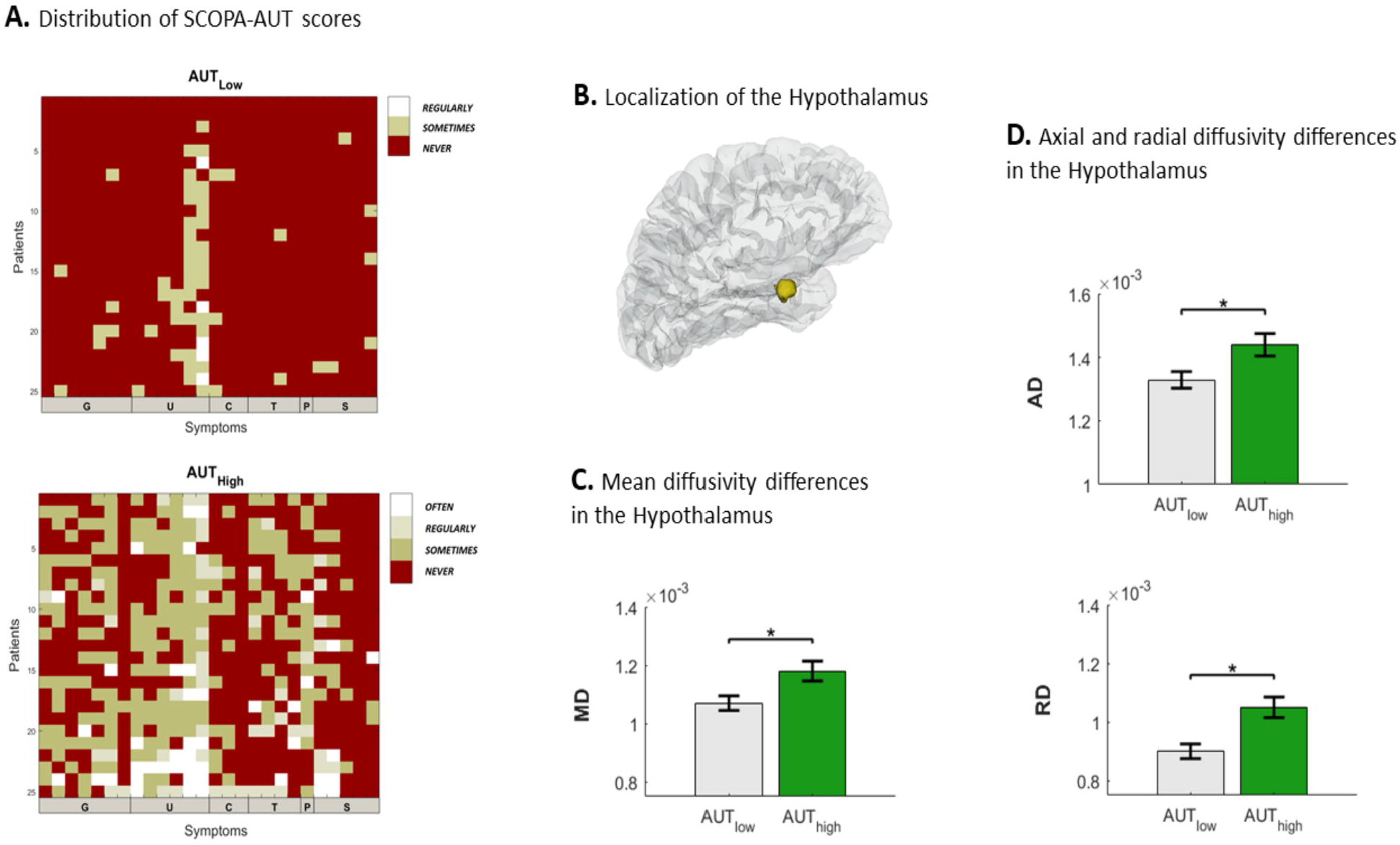
Hypothalamic diffusivity in patients with low and high autonomic symptom burden. (A.) SCOPA-AUT symptoms as reported by each individual subject in the AUT_low_ and AUT_high_ groups. Illustration is arranged according to questions addressing gastrointestinal (G; seven questions), urinary (U; six questions), cardiovascular (C; three questions), thermoregulatory (T; four questions), pupillomotor (P; one question) and sexual symptoms (S; three questions for men, two for women). (B.) Localization of the segmented Hypothalamus (yellow) in one representative Parkinson’s disease patient. (C-D) Significant differences (* p<0.050) were found in mean diffusivity (MD) axial diffusivity (AD) and radial diffusivity (RD) between patients with low (AUT_low_) and high (AUT_high_) autonomic symptom burden.

### Diffusivity properties

We extracted diffusivity measures from each patient’s segmented hypothalamus (Fig. 1B) to perform group comparisons (Table 2). The results reveal significantly decreased MD (t_48_ = - 2.578, p = 0.013; Fig. 1C), AD (t_48_ = −2.622, p = 0.012; Fig. 2D) and RD (t_48_ = −2.507, p = 0.016; Fig. 1D) in the AUT_low_ group, relative to the AUT_high_ group. While our main focus was on diffusivity measures, given earlier reports [22,23], we have additionally computed group differences in FA. We found a trend for stronger FA in the AUT_low_ group compared to the AUT_high_ group (p = 0.052).

**Figure 2:**
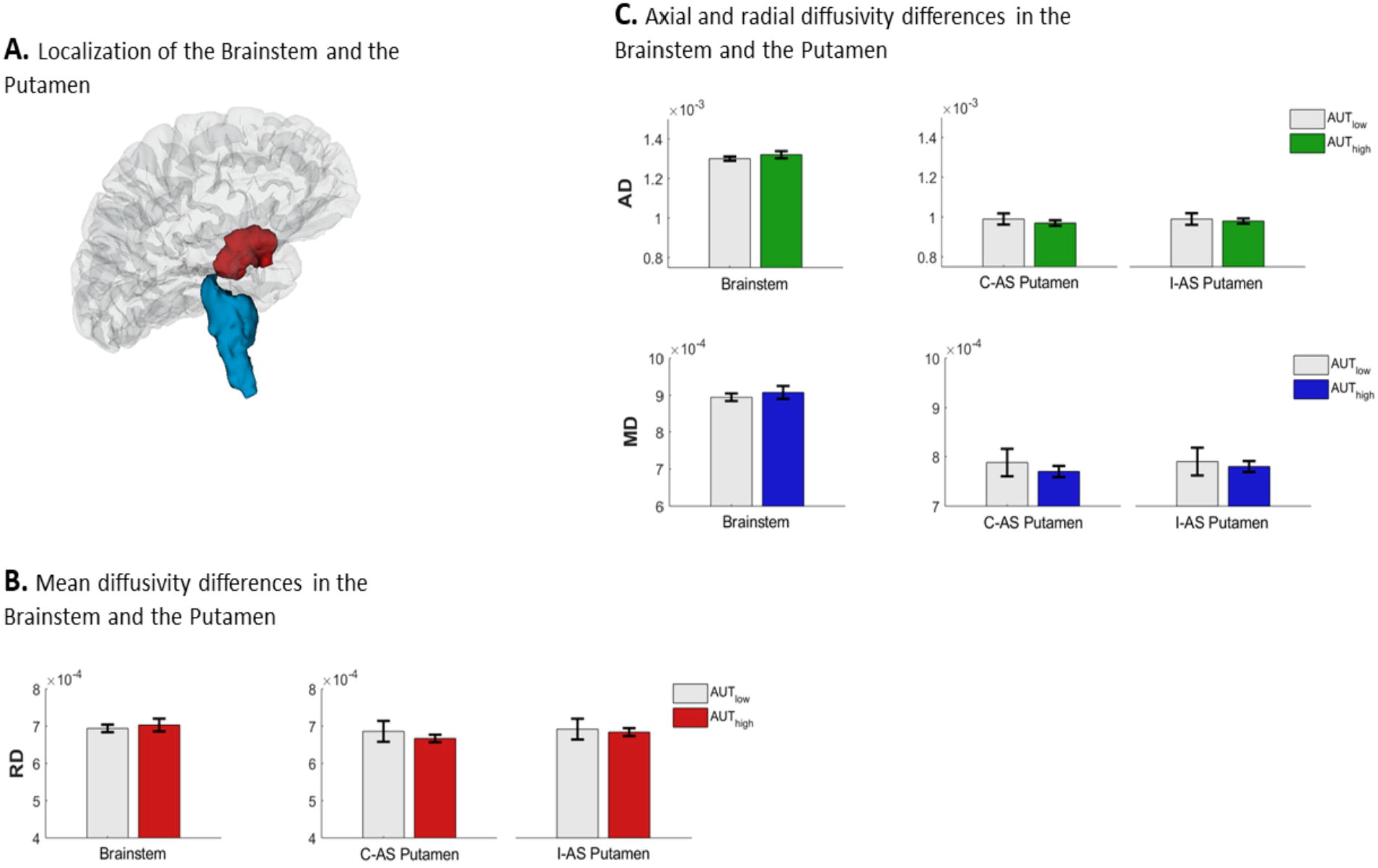
Diffusivity properties in the brainstem and the putamen. (A.) Localization of the segmented brainstem (blue) and Putamen (red), in one representative Parkinson’s disease patient. (B-C) No significant differences were found between patients with high (AUT_high_) and low (AUT_low_) autonomic symptom burden in mean, axial or radial diffusivity (MD, AD and RD), in the brainstem or in the putamen, contra (C-AS Putamen) and ipsilateral (I-AS Putamen) to the patients’ affected side.

**Table 2:** Diffusivity measures in the hypothalamus, the brainstem and the contralateral affected-side (AS) putamen of Parkinson’s disease patients in the low (AUT_low_) and high (AUT_high_) autonomic symptom burden groups

|  | *Hypothalamus* |  | *Brainstem* |  | *Contra AS Putamen* |  |
| --- | --- | --- | --- | --- | --- | --- |
| | $AUT_{low}$ | $AUT_{high}$ | $AUT_{low}$ | $AUT_{high}$ | $AUT_{low}$ | $AUT_{high}$ |
| **MD** ($\cdot 10^{-3}$) | $1.11 \pm 0.025$ | $1.18 \pm 0.034$ | $0.895 \pm 0.010$ | $0.909 \pm 0.017$ | $0.788 \pm 0.028$ | $0.770 \pm 0.011$ |
| **AD** ($\cdot 10^{-3}$) | $1.33 \pm 0.026$ | $1.44 \pm 0.035$ | $1.30 \pm 0.011$ | $1.32 \pm 0.018$ | $0.994 \pm 0.027$ | $0.975 \pm 0.013$ |
| **RD** ($\cdot 10^{-3}$) | $0.94 \pm 0.025$ | $1.05 \pm 0.034$ | $0.694 \pm 0.010$ | $0.703 \pm 0.017$ | $0.686 \pm 0.028$ | $0.667 \pm 0.011$ |

All values are mean ± standard errors. AD: Axial Diffusivity; MD: Mean Diffusivity; RD: Radial Diffusivity. Contra AS: contralateral to affected-side

Our comparison of the two groups’ demographic and clinical characteristics revealed significant group differences in age (Table 1). We next wished to ascertain that the differences in hypothalamic diffusivity were retained when controlling for this variable. Indeed, ANCOVA analyses revealed decreased diffusivity in the AUT_low_ group, relative to the AUT_high_ group, when age is controlled for (MD: F_1,48 =_ 4.459, p = 0.040; AD: F_1,48 =_ 4.181, p = 0.047; RD: F_1,48_ = 4.447, p = 0.040).

Our analysis demonstrates differences in hypothalamic diffusivity, between patients with low and high autonomic symptom burden. We next wished to establish the specificity of these differences to the hypothalamus. We considered two control ROIs. First, the brainstem, which comprises several structures implicated in autonomic function, notably the medulla oblongata, where diffusivity alterations were noted between Parkinson’s disease patients and healthy controls [15]. Diffusivity in the brainstem did not differ significantly between the two groups (MD: t_48_ = −0.676, p = 0.502; Fig. 2B; AD: t_48_ = −1.018, p = 0.314; RD: t_48_ = −0.475, p = 0.637; Fig. 2C).

A second control ROI we considered was the putamen, where alterations in multimodal MRI measures have been reported in Parkinson’s disease patients vs. controls [24,25]. There were no significant differences in putaminal diffusivity, neither ipsilateral (MD: t_48_ = 0.325, p = 0.747; AD: t_48_ = 0.432, p = 0.668; RD: t_48_ = 0.265, p = 0.792) nor contralateral (MD: t_48_ = 0.623, p = 0.536; AD: t_48_ = 0.603, p = 0.549; RD: t_48_ = 0.624, p = 0.536; Fig. 2C) to the patients’ affected side. These results thus suggest that the association between altered diffusivity measures in the hypothalamus and autonomic symptom burden in Parkinson’s disease is strongly specific.

## Discussion

We report that three common measures of diffusivity (MD, AD and RD), computed with diffusion MRI, were increased in patients with high autonomic symptom burden compared to patients with low burden. These alterations in hypothalamic diffusivity were retained when controlling for age, a factor that differed among the two groups of patients. Group differences in measures of diffusivity were specific to the hypothalamus, and were not observed in two control ROIs, the brainstem and the putamen.

As a key regulatory structure in the CAN, the hypothalamus has both afferent and efferent connections with other major structures in this network [9]. As such, the hypothalamus directly or indirectly participates in multiple autonomic functions, including homeostasis maintenance, processing of visceral sensory information [26], temperature control [27], and cardiovascular function [28]. Neurochemical abnormalities in the hypothalamus of patients with Parkinson’s disease were previously reported [9,10], along with Lewy body formations documented post-mortem [29]. A recent study demonstrated decreased functional connectivity between the hypothalamus and the striatum and thalamus in Parkinson’s disease patients with high autonomic symptom burden [14]. Our findings coincide with, and significantly add to, the literature by demonstrating that in addition to alterations in hypothalamic neurotransmitter levels [9], post-mortem examination [12,13], and functional connectivity [14], there appear to be measurable and significant differences in the in vivo microstructural properties of the hypothalamus among Parkinson’s disease patients with higher autonomic symptom burden.

The cellular mechanisms that underlie effects recorded using diffusion MRI are not fully understood. We found an increase in hypothalamic RD, in patients with higher autonomic symptom burden, which may reflect demyelination [30]. In addition, an increase in AD, as also observed in the current study, may be attributed to reduced axonal caliber [31]. These microstructural alterations may reflect reduced hypothalamic connectivity with other brain regions implicated in Parkinson’s disease neuropathology. These hypotheses should be investigated in future studies employing neuroanatomical, physiological, or neurochemical techniques, given that alterations in diffusivity measures may reflect multiple neuronal mechanisms [32].

The specificity of diffusivity changes in the hypothalamus with relation to autonomic symptoms in Parkinson’s disease demonstrated in this study, is clinically significant. Biomarkers in Parkinson’s disease in general, and in autonomic dysfunction in particular, are in urgent need [33]. Treatments for autonomic symptoms in Parkinsonian disorders are scarce, and lack specificity. Improved understanding of the underlying structural and functional neuronal mechanisms of Parkinson’s disease has produced excellent treatments for motor symptoms (namely, Levodopa); similarly, improved understanding of autonomic dysfunction in this disease may lead to the development of targeted therapies for these symptoms. Additionally, the identification of a disease marker specific to autonomic dysfunction, rather than simply related to overall disease severity, would greatly aid clinical trial efforts.

The diffusivity differences observed between patients with high and low autonomic symptom burden in the current study were specific to the hypothalamus. Similar differences were not seen in the brainstem or the putamen. The brainstem was chosen as a control ROI, given that some of its structures are key constituents in the CAN [7,8,34] and play key roles in moment-to-moment and reflexive control of autonomic functions. Diffusivity in the medulla oblongata has been previously shown to be increased in Parkinson’s disease patients relative to controls, and is correlated with heart rate and respiratory frequency variability recorded during rapid eye movement sleep [35]. We did not find diffusivity differences in the brainstem between Parkinson’s disease patients with high and low burden of autonomic symptoms. However, the study by Pyatigorskaya and colleagues [35] focused on cardiovascular and respiratory functions, comparing Parkinson’s disease patients and controls, rather than autonomic symptom burden as studied here. The study also included Parkinson’s disease patients with a wider range of disease durations, thus a comparison with the current results is challenging, as our cohort was more uniform in that respect. It is noteworthy however, that while the medulla oblongata has projections to the hypothalamus [9,36], hypothalamic outputs to this structure have also been identified [9,37]. Thus, a possibility that remains to be tested in future longitudinal studies is that diffusivity alterations in the medulla oblongata and the hypothalamus emerge at different disease stages [38]. Such longitudinal investigations could also compare the sensitivity, specificity and predictive power of diffusivity measures extracted from different CAN structures in tracking the progression of autonomic symptoms in Parkinson’s disease. This may be particularly important in the development of clinical biomarkers for autonomic dysfunction in this disease.

A second control region we have tested and where no diffusivity differences were found between the two groups was the putamen, both ipsi- and contralateral to the patients’ predominantly affected side. Putaminal diffusivity alterations have been reported in Parkinson’s disease patients when compared to controls [39]. Diffusivity differences in the putamen were also consistently found [40] when comparing Parkinson’s disease patients to patients with multiple systems atrophy (MSA), particularly those with the Parkinsonian variant of the disease (MSA-P). Though MSA patients frequently exhibit autonomic dysfunction, and involvement of the putamen in autonomic dysfunction in parkinsonism may be postulated, this is not supported by the current results. Studies specifically comparing hypothalamic and putaminal diffusivity in MSA and Parkinson’s disease patients with dysautonomia are needed in order to more closely delineate the role of the putamen in autonomic dysfunction in these conditions.

A few potential limitations need to be considered. First, the relative homogeneity of the population examined in this study must be considered. Though in many ways this is a strength of the current study, future studies will be necessary to investigate microstructural changes and functional connectivity of CAN structures in a more diverse Parkinson’s disease population. Additionally, autonomic symptom burden in the current study was only based on a screening questionnaire (the SCOPA-AUT [18]), which may be less accurate and specific than quantitative physiologic autonomic testing. Thus, although this questionnaire is considered a reliable instrument [19], investigating microstructural alterations within and outside the hypothalamus when using quantitative autonomic testing is warranted. Finally, while 3T MRI provides sufficient resolution for investigations of supratentorial structures, the brainstem is somewhat difficult to delineate with this method. Future studies with higher resolution imaging modalities, such as 7T MRI, may allow to better distinguish diffusivity in the medulla or medullary substructures from other brainstem structures.

In summary, the present study highlights altered microstructural properties in the hypothalamus, seen in Parkinson’s disease patients with high as compared to low autonomic symptom burden. These findings may lead to the development of an in vivo biomarker that can monitor autonomic dysfunction in Parkinson’s disease and be used in clinical trials testing pharmacological or other therapeutic interventions.

## Acknowledgements

Data were obtained from the PPMI database. For up-to-date information on the study, visit www.ppmi-info.org. PPMI, a public-private partnership, is funded by the Michael J.Fox Foundation for Parkinson’s Research and funding partners, including Abbvie, Avid Radiopharmaceuticals, Biogen, Bristol-Myers Squibb, Covance, GE healthcare, Genentech, GlaxoSmithKline, Lilly, Lundbeck, Merck, Meso Scale Discovery, Pfizer, Piramal, Roche, Servier, and UCB. Additional funding was provided by NIH grant U54HD079124.

## Author Contributions

M.Sk, N.B, Y.L and E.D contributed to the conception and design of the study. C.R, J.P, A.B, M.St and E.D contributed to data analysis. C.R, M.Sk, N.B and E.D contributed to drafting a significant portion of the manuscript. All authors read and approved the manuscript.

